# Cognitive ability, Socioeconomic Status, and Depressive Symptoms: A Gene-Environment-Trait Correlation

**DOI:** 10.1101/727552

**Authors:** Reut Avinun

## Abstract

**Purpose:** Depression is genetically influenced, but the mechanisms that underlie these influences are largely unknown. Recently, shared genetic influences were found between depression and both cognitive ability and educational attainment (EA). Although genetic influences are often thought to represent direct biological pathways, they can also reflect indirect pathways, including modifiable environmental mediations (gene-environment-trait correlations). Here, I tested whether the genetic correlation between cognitive ability and depressive symptoms partly reflects an environmental mediation involving socioeconomic status (SES).

**Methods:** As previously done to increase statistical power, and due to their high phenotypic and genetic correlation, EA was used as a proxy for cognitive ability. Summary statistics from a recent genome-wide association study of EA were used to calculate EA polygenic scores. Two independent samples were used: 522 non-Hispanic Caucasian university students from the Duke Neurogenetics Study (277 women, mean age 19.78±1.24 years) and 5,243 white British volunteers (2,669 women, mean age 62.30±7.41 years) from the UK biobank.

**Results:** Mediation analyses in the two samples indicated that higher proxy-cognitive ability polygenic scores predicted higher SES, which in turn predicted lower depressive symptoms.

**Conclusion:** Current findings suggest that some of the genetic correlates of depressive symptoms depend on an environmental mediation and consequently that modifying the environment, specifically through social and economic policies, can affect the genetic influences on depression. Additionally, these results suggest that findings from genetic association studies of depression may be context-contingent and reflect social, cultural, and economic processes in the examined population.

---

Depression is a major cause of disability. It has a global prevalence of around 4.7% [1], and it is predicted to become one of the three leading causes of illness by 2030 [2]. Interestingly, low cognitive ability in childhood has been shown to predict high levels of depression (e.g., [3,4]). Educational attainment (EA), which is often used as a proxy for cognitive ability, has been also linked to depression, so that the probability of experiencing depression decreases for additional years of education [5]. Recent studies have found negative genetic associations between depression and both cognitive ability [6] and EA [7]. Furthermore, by employing a genetically informed analysis (Mendelian randomization), both relatively low cognitive ability and low EA were marked as risk factors for depression [8,7]. Notably, how the genetic correlation between cognitive ability and depression is mediated has not been established.

Recently, it was hypothesized that socioeconomic status (SES) may mediate the association between cognitive ability and depression [9]. More generally, it was suggested that the environment may mediate genetic correlations between two phenotypes within the same individual, in a process termed gene-environment-trait correlations [9]. This hypothesis stems from accumulating research showing passive, active, and evocative processes that lead to correlations between genetic variation and environmental measures, such as parenting and stressful life events [10,11]. These passive, active, and evocative processes, known as gene-environment correlations [12,13], occur due to genetically influenced characteristics that shape the individual’s environment. As the environment can in turn substantially affect various outcomes, it may act as a mediator of genetic effects within the same individual and contribute to the widespread genetic correlations observed between numerous phenotypes [14], including cognitive ability and depression [8,6]. In other words, it is possible that genetic influences on two different phenotypes, like cognitive ability and depression, are linked, because one genetically influenced phenotype (e.g., cognitive ability) affects an environment (e.g., SES) that, in turn, affects another phenotype (e.g., depressive symptoms).

Identifying gene-environment-depression correlations can shed light on environments that play a role in pathways that connect between certain genetic variations and depression. Disrupting such pathways through public policy will modify these indirect genetic influences. Additionally, such environmental mediations can demonstrate the importance of context in the discovery of the genetic correlates of depression, because different contexts can translate into different environmental mediations of genetic effects.

SES, which can be defined as an individual’s or group’s position within a social hierarchy that is determined by factors such as education, occupation, income, and wealth [15], has been shown to be genetically influenced [16,17]. Put differently, genetically influenced traits affect an individual’s SES. One of these traits, as has been found in a meta-analysis of longitudinal studies, is cognitive ability [18], which is highly heritable [19]. Because SES has been associated with various physiological and mental disorders (e.g., [15,20,21]), including depression [22], and a genetic correlation between SES and depression has also been observed [16], a gene-environment-trait correlation in which SES mediates the genetic correlation between cognitive ability and depression, is possible.

Sample sizes of more than a million individuals are needed for reliable detection of relevant genetic variation in genome wide association studies (GWASs) of complex traits such as cognitive ability [23]. Because such sample sizes with assessments on cognitive ability are challenging to obtain, it is common to use EA as a proxy for cognitive ability (e.g., [24,25]) to increase statistical power. Other than their high phenotypic correlation, there is also a high genetic correlation between EA and cognitive ability, indicating shared genetic influences (a single nucleotide polymorphism-based genetic correlation of .95; [17]). A recent GWAS of EA [26] included ~1.1 million European-descent participants, making it one of the most powerful, and consequently prevalently used, GWASs in psychology (for comparison, a recent GWAS of cognitive ability included 269,867 individuals; [6]). A polygenic score based on the summary statistics from this GWAS explained ~11% of the variance in EA. In the current study, I tested whether SES mediated an association between EA polygenic scores, used as a proxy for cognitive ability polygenic scores, and depressive symptoms.

Two independent samples were used for the analyses: a sample of 522 non-Hispanic Caucasian university students from the Duke Neurogenetics Study and a sample of 5,243 adult white British volunteers from the UK Biobank (UKB). Notably, the UK biobank is the main sample in the cognitive ability GWAS (195,653 of the 269,867 individuals; [6]), which consequently also favors the use of the EA GWAS in the current study. The DNS and the UKB complement each other in several ways: 1) the measures used for the assessment of SES and depressive symptoms differed in the two samples as will be detailed below, and therefore finding a significant mediation in both samples would suggest that the result is robust to different operationalizations of these two constructs; 2) the two samples represented different age groups, young adulthood and older adulthood, and therefore finding the hypothesized mediation in both samples could show that it is not specific to a particular age range; 3) as it can be argued that EA is a measure of SES, in the DNS all participants were students at the same university, which is similar to controlling for EA in this sample; and 4) in the UKB it was possible to test a longitudinal mediation model, which can provide further support for causal inference. Lastly, as the EA GWAS included data from the UKB, in the analyses of the UKB data EA polygenic scores were based on summary statistics from a GWAS that did not include the UKB as a discovery sample (obtained from Dr. Aysu Okbay, who is one of the authors of the original GWAS).

## Materials and Methods

### Participants

The Duke Neurogenetics Study (DNS) sample consisted of 522 self-reported non-Hispanic Caucasian participants (277 women, mean age 19.78±1.24 years) who were not related and for whom there was complete data on genotypes, SES, depressive symptoms, and all covariates. Participants were recruited through posted flyers on the Duke University campus and through a Duke University listserv. All procedures were approved by the Institutional Review Board of the Duke University Medical Center, and participants provided informed consent before study initiation. All participants were free of the following study exclusions: 1) medical diagnoses of cancer, stroke, diabetes requiring insulin treatment, chronic kidney or liver disease, or lifetime history of psychotic symptoms; 2) use of psychotropic, glucocorticoid, or hypolipidemic medication; and 3) conditions affecting cerebral blood flow and metabolism (e.g., hypertension).

The UKB sample consisted of 5,243 white British volunteers (2,669 women, mean age 62.30±7.41 years), who participated in the UKB’s first assessment and the imaging wave, completed an online mental health questionnaire [27], and had complete genotype, SES, depressive symptoms and covariate data. The UKB (www.ukbiobank.ac.uk; [28]) includes over 500,000 participants, between the ages of 40 and 69 years, who were recruited within the UK between 2006 and 2010. The UKB study was approved by the National Health Service Research Ethics Service (reference: 11/NW/0382), and current analyses were conducted under UKB application 28174 (because the application originally included a request for neuroimaging data, the sample used in this study is limited to individuals who participated in the imaging wave).

### Ancestry

Because self-reported race and ethnicity are not always an accurate reflection of genetic ancestry, an analysis of identity by state of whole-genome SNPs in the DNS was performed in PLINK v1.9 [29]. Before running the multidimensional scaling (MDS) components analysis, SNPs were pruned for high LD (r^2^>0.1), and the following were removed: C/G and A/T SNPs, SNPs with a missing rate >.05 or a minor allele frequency <.01, SNPs that did not pass the Hardy-Weinberg equilibrium test (p<1e-6), sex chromosomes, and regions with long range LD (the MHC and 23 additional regions; [30]). The first two MDS components computed for the non-Hispanic Caucasian subgroup, as determined by both self-reports and the MDS components of the entire mixed race/ethnicity DNS sample, were used as covariates in analyses of data from the DNS. The decision to use only the first two MDS components was based on an examination of a scree plot of eigenvalues, which became very similar after the second MDS component (additional information and plots are available at https://www.haririlab.com/methods/genetics.html).

For analyses of data from the UKB, only those who were ‘white British’ based on both self-identification and a genetic principal components analysis were included. Additionally, the first 10 principal components received from the UKB’s data repository (unique data identifiers: 22009.0.1-22009.0.10) were included as covariates as previously done (e.g., [31,32]). Further details on the computation of the principal components can be found elsewhere (http://www.ukbiobank.ac.uk/wp-content/uploads/2014/04/UKBiobank_genotyping_QC_documentation-web.pdf).

### Socioeconomic status

In the DNS, SES was assessed using the “social ladder” instrument [33], which asks participants to rank themselves relative to other people in the United States (or their origin country) on a scale from 0–10, with people who are best off in terms of money, education, and respected jobs, at the top (10) and people who are worst off at the bottom (0).

In the UKB, SES was assessed based on the report of average household income before tax, coded as: 1 - Less than 18,000; 2 - 18,000 to 31,000; 3 - 31,000 to 52,000; 4 - 52,000 to 100,000; and 5 - Greater than 100,000. The reports made during the first assessment (i.e., before the evaluation of depressive symptoms), between 2006 and 2010, were used.

### Depressive symptoms

In the DNS, the 20-item Center for Epidemiologic Studies Depression Scale (CES-D) was used to assess depressive symptoms in the past week [34]. All items were summed to create a total depressive symptoms score.

In the UKB, the Patient Health Questionnaire 9-question version (PHQ-9) was used to assess depressive symptoms in the past 2 weeks [35]. The participants answered these questions during a follow-up between 2016 and 2017. All items were summed to create a total depressive symptoms score.

### Genotyping

In the DNS, DNA was isolated from saliva using Oragene DNA self-collection kits (DNA Genotek) customized for 23andMe (www.23andme.com). DNA extraction and genotyping were performed through 23andMe by the National Genetics Institute (NGI), a CLIA-certified clinical laboratory and subsidiary of Laboratory Corporation of America. One of two different Illumina arrays with custom content was used to provide genome-wide SNP data, the HumanOmniExpress (N=328) or HumanOmniExpress-24 (N=194; [36–38]). In the UKB, samples were genotyped using either the UK BiLEVE (N=501) or the UKB axiom (N=4,742) array. Details regarding the UKB’s quality control can be found elsewhere [39].

### Quality control and polygenic scoring

For genetic data from both the DNS and UKB, PLINK v1.90 [29] was used to apply quality control cutoffs and exclude SNPs or individuals based on the following criteria: missing genotype rate per individual >.10, missing rate per SNP >.10, minor allele frequency <.01, and Hardy-Weinberg equilibrium p<1e-6. Additionally, in the UKB, quality control variables that were provided with the dataset were used to exclude participants based on a sex mismatch (genetic sex different from reported sex), a genetic relationship to another participant, outliers for heterozygosity or missingness, and UKBiLEVE genotype quality control for samples (unique Data Identifiers 22010.0.0, 22011.0.0-22011.0.2, 22018.0.0, 22050.0.0-22052.0.0).

Polygenic scores were calculated using PLINK’s [29] “--score” command based on published SNP-level summary statistics from the most recent EA GWAS [26]. Published summary statistics do not include the data from 23andMe per the requirements of this company (i.e., the sample of the GWAS the summary statistics for the DNS relied on included about 766,345 individuals). For the UKB analyses, summary scores from a GWAS that did not include the UKB as a discovery sample were used (i.e., the sample of the GWAS the summary statistics for the UKB relied on included about 324,162 individuals). SNPs from the GWAS of EA were matched with SNPs from the DNS and the UKB and for each SNP the number of the alleles (0, 1, or 2) associated with EA was multiplied by the effect estimated in the GWAS. The polygenic score for each individual was an average of weighted EA-associated alleles. This EA polygenic score was used as a proxy for cognitive ability genetic correlates. All SNPs matched with genotyped SNPs from the DNS and UKB were used regardless of effect size and significance in the original GWAS, as previously recommended and shown to be effective [40,41].

### Statistical analysis

The PROCESS SPSS macro, version 3.1 [42], was used to conduct the mediation analyses in SPSS version 26. Participants’ sex (coded as 0=males, 1=females), age, and ancestry (two genetic components for the DNS and 10 for the UK biobank) were entered as covariates in all analyses. Bias-corrected bootstrapping (set to 5,000) was used in the mediation analyses to allow for non-symmetric 95% confidence intervals (CIs). Specifically, indirect effects are likely to have a non-normal distribution, and consequently the use of non-symmetric CIs for the determination of significance is recommended [43]. To complement the bias-corrected bootstrapping method and add supportive evidence for the indirect effect [44], I also present the test of joint significance, which examines whether the *a path* (proxy-cognitive ability polygenic scores to SES) and the *b path* (SES to depressive symptoms, while controlling for the proxy-cognitive ability polygenic scores) are significant. The proxy-cognitive ability polygenic scores were standardized (i.e., M=0, SD=1) in SPSS to make interpretability easier.

As a post-hoc analysis, in the UKB it was possible to analyze the longitudinal data while excluding those who reported on ever seeing a general physician (N=1,843) or a psychiatrist (N=501) “for nerves, anxiety, tension or depression”, at the first assessment (i.e., at the first assessment both household income and these two questions regarding the experience of depression were reported, and more than 6 years later information on depressive symptoms, as assessed by the PHQ-9, was collected). By excluding participants who experienced depression before the assessment of household income (i.e., SES), a significant prediction of later depressive symptoms is more likely to be causal.

## Results

### Descriptive statistics

In the DNS, the SES measure ranged between 2 and 10 (M=7.34, SD=1.43) and depressive symptoms ranged between 0 and 43 (M=8.94, SD=7.13). In the UKB, the SES measure ranged between 1 and 5 (M=2.92, SD=1.11), and depressive symptoms, estimated about 6 years later, ranged between 0 and 27 (M=2.50, SD=3.43).

### Proxy-cognitive ability polygenic scores and SES (a path) in the DNS

The proxy-cognitive ability polygenic scores were significantly associated with SES (b=.20, SE=.06, p=.0016; R^2^=0.018), so that higher scores predicted higher SES. Of the covariates, age and sex were significantly associated with SES, so that older participants (b=.13, SE=.05, p=.008) and men (b=−.45, SE=.12, p=.0003) were characterized by higher SES.

### SES and depressive symptoms (b path) in the DNS

With the proxy-cognitive ability polygenic scores in the model, SES significantly and negatively predicted depressive symptoms (b=−.61, SE=.22, p=.007; R^2^=0.014). Higher SES predicted lower depressive symptoms. Of the covariates, age was significantly associated with depressive symptoms, so that being younger was associated with higher depressive symptoms (b=−.53, SE=.25, p=.037).

### Proxy-cognitive ability polygenic scores and depressive symptoms in the DNS

The proxy-cognitive ability polygenic scores did not significantly predict depressive symptoms (b=−.11, SE=.32, p=.74). Notably, however, the significance of a direct path from X (proxy-cognitive ability polygenic scores) to Y (depressive symptoms) or the ‘total effect’ (the ‘c’ path), is not a prerequisite for testing a mediation/indirect effect [45–47], which was the main aim of the current study.

### Indirect Effects in the DNS

The indirect path (*a*\**b*), proxy-cognitive ability polygenic scores to depressive symptoms via SES, was significant as indicated by the bias corrected bootstrapped 95% CI not including zero (Figure 1a; indirect effect=−.12, bootstrapped SE=.06, bootstrapped 95% CI: −.26 to −.02). The indirect effect remained significant when 10 MDS components of genetic ancestry (instead of the initial 2) and genotyping platform were included as covariates (i.e., in addition to sex and age; indirect effect=−.12, bootstrapped SE=.06, bootstrapped 95% CI: −.27 to −.02).

**Figure 1.**
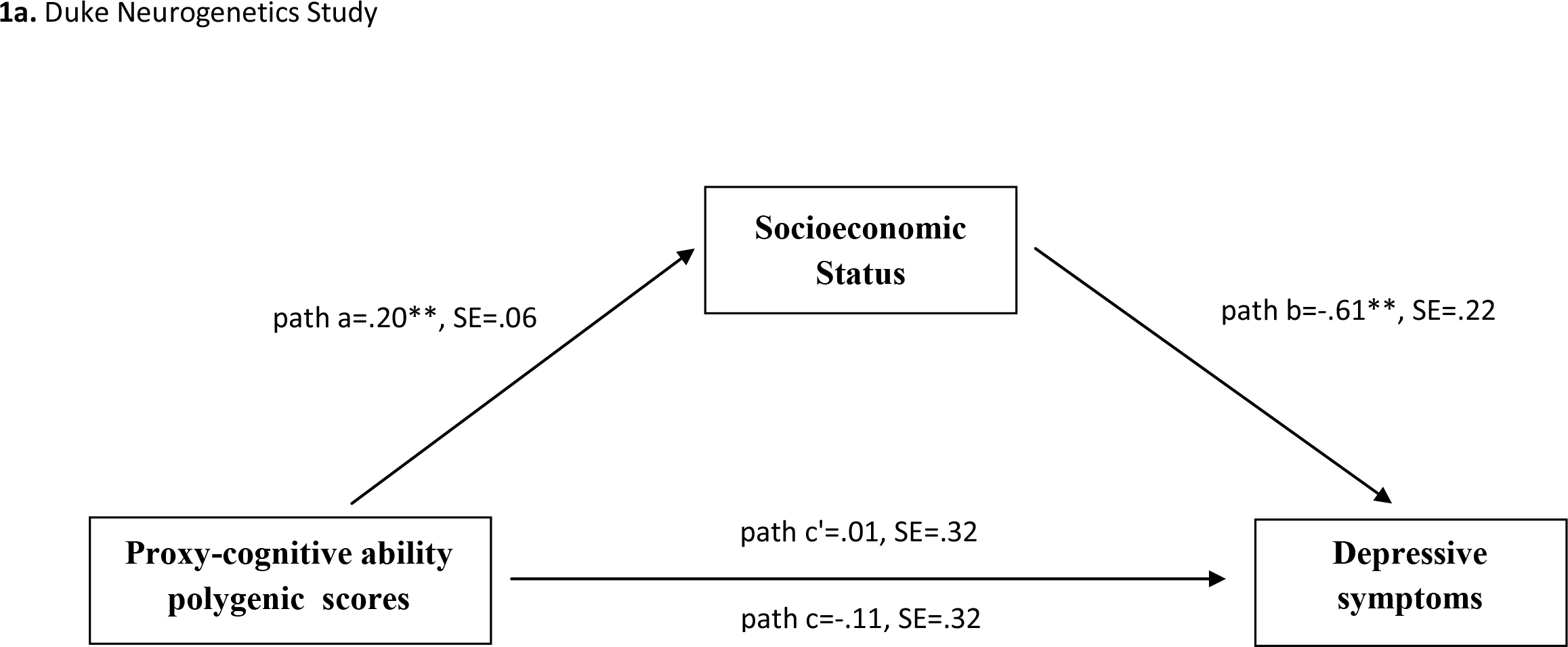
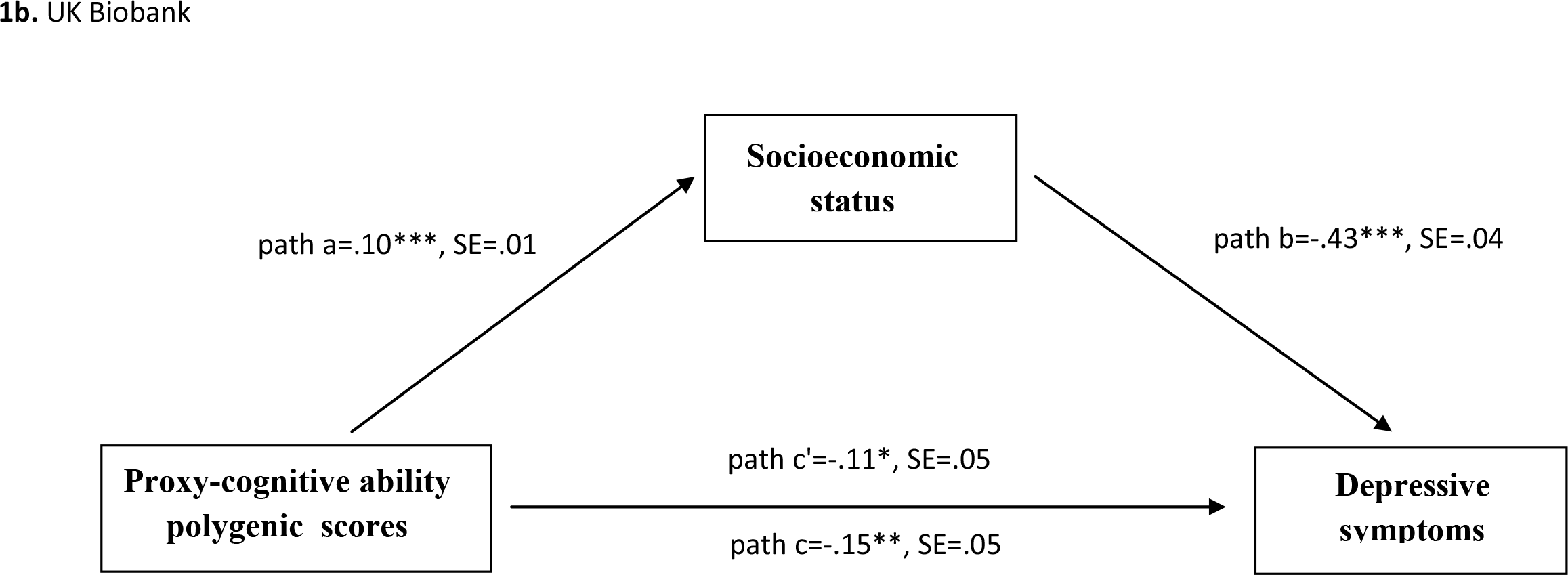
Mediation model linking genetic influences on cognitive ability to depressive symptoms, via socioeconomic status. *Note*. *p<.05, **p<.01, ***p<.0001. c- the total effect of the proxy-cognitive ability polygenic scores on depressive symptoms; c’-the effect of proxy-cognitive ability polygenic scores on depressive symptoms, while controlling for socioeconomic status.

### Proxy-cognitive ability polygenic scores and SES (a path) in the UKB

For the UKB, proxy-cognitive ability polygenic scores that were based on a GWAS that did not include the UKB as a discovery sample, were used. Additionally, genotyping platform was included as a covariate in the analyses. The proxy-cognitive ability polygenic scores were significantly associated with SES (b=.10, SE=.01, p<.0001, R^2^=0.008), indicating that higher scores predicted higher SES. Of the covariates, age and sex were significantly associated with SES. Younger participants (b=−.05, SE=.002, p<.0001) and men (b=−.24, SE=.03, p<.0001) were characterized by higher SES.

### SES and depressive symptoms (b path) in the UKB

With the proxy-cognitive ability polygenic scores in the model, SES significantly and negatively predicted depressive symptoms (b=−.43, SE=.04, p<.0001, R^2^=0.017), so that higher SES predicted lower depressive symptoms. The covariates age and sex were significantly associated with depressive symptoms, revealing that younger ages (b=−.10, SE=.007, p<.0001) and being a woman (b=.48, SE=.09, p<.0001) were associated with higher depressive symptoms.

### Proxy-cognitive ability polygenic scores and depressive symptoms in the UKB

Higher proxy-cognitive ability polygenic scores were significantly associated with lower depressive symptoms (b=−.15, SE=.05, p=.001).

### Indirect effect in the UKB

The indirect path was significant (Figure 1b; indirect effect=−.04, bootstrapped SE=.008, bootstrapped 95% CI: −.06 to −.03). To test the robustness of the finding, a post-hoc analysis that excluded participants who, at the first assessment, reported on ever seeing a professional for nerves or depression (leaving 3,447 participants), was conducted. This was done in an attempt to increase the likelihood of only including the individuals who became depressed between the first assessment, when household income was first reported, and the assessment more than 6 years later, in which depressive symptoms were assessed. Notably, a correlation between the report of household income at the first assessment and the report of household income at the second assessment indicated some change in household income during this time period (r(5,243)=.68). Indeed, the longitudinal analysis supported a causal mediation, in which higher proxy-cognitive ability polygenic scores predicted higher SES, which in turn predicted lower depressive symptoms (*a path*: b=.08, SE=.02, p<.0001, R^2^=0.005; *b path*: b=−.15, SE=.04, p=.0003, R^2^=0.004; indirect effect=−.012, bootstrapped SE=.004, bootstrapped 95% CI: −.022 to −.005).

## Discussion

The current results suggest that the negative genetic correlation previously observed between cognitive ability and depressive symptoms is partly mediated by SES, an environment that can be modified through social and economic policies. The indirect effect was found in two independent samples with different characteristics and measures, demonstrating the robustness of the associations. Notably, in the UKB the indirect effect was tested longitudinally, with data on SES that was collected about 6 years before the assessment of depressive symptoms. A supplementary analysis that excluded participants who reported ever seeing a professional for nerves or depression at the first assessment, was also significant, further supporting a causal temporal mediation by predicting change in depressive symptoms.

The found mediation supports the gene-environment-trait correlations hypothesis (rGET; [9]), which suggests that certain genetic correlations between different phenotypes may be mediated, at least in part, by the environment, i.e., an environmentally mediated pleiotropy. The found proxy-cognitive ability polygenic scores→SES→depressive symptoms mediation stresses the importance of context in genetic studies of depression. The genetic correlates of depression in one population may be different from the ones found in another population. Put differently, because the environment can act as a mediator of some of the genetic influences on depression, if the environment differs between GWASs, the captured genetic influences will differ. Importantly, the current results suggest that social policies aimed at reducing socioeconomic inequalities may weaken the genetic effects on depression by disrupting the pathway that leads to the association between cognitive ability and depression.

Low SES may be a risk factor for depression by leading to an increase in life stress that stems from having to make ends meet and from living in a disadvantaged neighborhood, which is associated with higher crime and fewer resources [48]. Low SES has also been associated with poorer access to green spaces [49], and with health damaging behaviors, such as physical inactivity, higher alcohol consumption, and poor nutrition [50,51], which are thought to affect mental health (e.g., [52,32,53]). All of these mediators can be possible targets for policy makers.

The strengths of the current study include the use of two independent samples with markedly different measures and characteristics (e.g., young university students versus older community volunteers) and a GWAS-derived polygenic score, but it is also limited in ways that can be addressed in future studies. First, the findings are limited to populations of European descent and to the Western culture. Second, both samples consisted of volunteers and consequently do not fully represent the general population. However, it may be speculated that the observed associations would strengthen with the inclusion of more individuals from low SES backgrounds, which are usually characterized by higher levels of depression [54]. Third, the mediation model should be replicated within longitudinal designs in which the same measures of SES and depressive symptoms are available at multiple time points.

In conclusion, the current results shed light on the genetic associations that have been observed between cognitive ability and depression [6], and suggest that they are partly mediated by SES. The mediation by SES is important because it suggests that the genetic influences on depression may be moderated by public policy and that the genetic composition of depression depends on the social context in which it is examined.

## Acknowledgements

I would like to thank the participants of the Duke Neurogenetics Study and the members of the Laboratory of NeuroGenetics, especially Annchen R. Knodt, Spenser R. Radtke, and Bartholomew D. Brigidi for their assistance with data collection and analysis. I would also like to thank the head of the laboratory, Prof. Ahmad Hariri, without whom this study would not have been possible. Lastly, I would like to thank Dr. Aysu Okbay for her help with obtaining a GWAS of educational attainment that did not include UK biobank data.

## Availability of data and material

The required procedures for obtaining the DNS data are detailed on our website https://www.haririlab.com/projects/procedures.html. The UK Biobank data requires contacting the UK Biobank team directly, through http://www.ukbiobank.ac.uk.

## Code availability

Available from the author.

